## Supplementary tables for "Entorhinal layer 6b subplate neurons govern spatial learning and memory"

Supplementary Table 1: Summary of pSPN membrane properties.

| **Property**^1,2^ | **Mean** | **SEM** |
| --- | --- | --- |
| Resting membrane potential (mV) | -68.7345 | 1.18683 |
| AP threshold potential (mV) | -48.1541 | 1.39943 |
| AP peak^3^ (mV) | 83.34582 | 2.15426 |
| AP rise time^4^ (ms) | 0.605888 | 0.04134 |
| AP width^5^ (ms) | 1.513645 | 0.04891 |
| Max rise slope | 230.1616 | 9.25946 |
| Max decay slope | 137.6243 | 4.31897 |
| Afterhyperpolarization^6^ (mV) | 11.54226 | 1.40138 |
| Latency to first AP (ms) | 303.2677 | 38.3723 |
| Input resistance (MΩ) | 487.4063 | 23.1846 |
| Sag ratio^7^ (%) | 5.8414 | 1.281 |

1. N=31 cells.
2. For each cells, all AP properties were measured for the first AP at the rheobase current.
3. Measured from 0.
4. 20%-80% of peak.
5. Full-width at half-maximum.
6. Relative to the threshold potential.
7. Measured as the ratio between the membrane potential at the positive peak at the start of the response to a -100 pA current injection, 1 s long, in relation to the average potential at the final 100 ms before the end of the pulse.

Supplementary Table 2: Summary of plasmids.

| Plasmid name | Source (deposited by) | Cat# |
| --- | --- | --- |
| pCAG-B19N | AddGene (I. Wickersham) | #59924 |
| pCAG-B19P | AddGene (I. Wickersham) | #59925 |
| pCAG-B19L | AddGene (I. Wickersham) | #59922 |
| pAdDeltaF6 | AddGene (J. Wilson) | #112867 |
| rAAV-DJ RepCap | Gift from Mark A. Kay |  |
| rAAV2-retro helper | AddGene (A. Karpova and D. Schaffer) | #81070 |
| pAAV-DIO-Ef1a-TVA-2A-N2cG | AddGene (Y. Ben Simon and P. Jonas) | #172360 |
| pAAV-EF1a-Cre | AddGene (K. Deisseroth) | #55636 |
| pAAV-SEO-CaMKIIa-TVA-2A-N2cG | AddGene (Y. Ben Simon and P. Jonas) | #172363 |
| pAAV-FRT-Ef1a-TVA-2A-N2cG | AddGene (Y. Ben Simon and P. Jonas) | #172361 |
| pAAV-SEO-CaMKII-EGFP | AddGene (Y. Ben Simon and P. Jonas) | #172361 |
| pAAV-DIO-hSyn-mCherry | AddGene (K. Deisseroth) | #114472 |
| RVdG-CVS-N2c-EGFP | AddGene (T. Jessell) | #73461 |
| RVdG-CVS-N2c-tdTomato | AddGene (T. Jessell) | #73462 |
| RVdG-CVS-N2c-tdTomato-ChIEF | AddGene (Y. Ben Simon and P. Jonas) | #172370 |
| RVdG-CVS-N2c-nl.EGFP-SypGEFP | AddGene (Y. Ben Simon and P. Jonas) | #172380 |

Supplementary Table 3: Summary of transgenic lines.

| Transgenic line | Full name | Source (deposited by) | Cat# |
| --- | --- | --- | --- |
| Ai9 | B6.Cg-Gt(ROSA)26Sor^tm9(CAG-tdTomato)Hze^/J | Jackson labs (H. Zeng) | 007909 |
| GAD1-EGFP | B6.Cg-Tg(Gad1-EGFP)3Gfng/J | Jackson labs (J. Huang) | 007677 |
| RCE-FRT | Gt(ROSA)26Sor^tm1.2(CAG-EGFP)Fsh^/Mmjax | Jackson labs (G. Fishell) | 32038 |
| Prox1-cre | Tg(Prox1-cre)SJ32Gsat/Mmucd | MMRRC (N. Heintz) | 036644-UCD |
| KA1-cre | C57BL/6-Tg(Grik4-cre)G32-4Stl/J | Jackson labs (S. Tonegawa) | 006474 |
| Dlx5/6-FlpE | Tg(mI56i-flpe)39Fsh/J | Jackson labs (G. Fishell) | 010815 |
| Ctgf-2A-dgCre | B6.Cg-Ccn2^tm1.1(folA/cre)Hze/^J | Jackson labs (H. Zeng) | 028535 |
| Ai32 | B6.Cg-Gt(ROSA)26Sor^tm32(CAG-COP4*H134R/EYFP)Hze^/J | Jackson labs (H. Zeng) | 024109 |
| Ai40 | B6.Cg-Gt(ROSA)26Sor^tm40.1(CAG-aop3/EGFP)Hze^/J | Jackson labs (H. Zeng) | 021188 |
| RosaDTA | B6.129S6(Cg)-Gt(ROSA)26Sor^tm1(DTA)Jpmb^/J | Jackson labs (M. Capecchi) | 010527 |
